# The risks of chronic exposure to ultraviolet light in laboratory environment

**DOI:** 10.1101/2023.12.30.572924

**Authors:** Eric Almeida Xavier

## Abstract

The use of ultraviolet (UV) sources in laboratories requires safety practices. For example, when the ultraviolet light of laminar flow is on, it is important that the researcher or team members do not remain in the environment. There are laminar flows that have glasses with an ultraviolet light filter, but many models do not have any protection against ultraviolet light. Thus, we measured the irradiance inside and outside two types of laminar flows without UV filter and with UV filter respectively. And calculated the radiation levels that a researcher could receive at one meter distance for one hour daily in a time interval of one year. Thus, we found that chronic exposure to UV radiation from laminar flow can cause health risks. And we concluded that all laboratories must adopt safety measures, for example does not allow the presence of people during the disinfection stage of laminar flow with UV light.

## Introduction

Currently, there are no workplace related rules and regulations set by Occupational Safety and Health Association (OSHA) in regard to UVC environmental health and safety. [1]. Like this there are many applications for ultraviolet field lights, for example environmental decontamination, water treatment and medical applications [2.3,4].

Thus, ultraviolet (UV) radiation is known to play a significant role in the development of cancer in skin, because of this, exposure precautions are recommended when using UVC light. [5]. For example, a dose of approximately 8 J/cm2, which corresponds to the UVA dose received approximately within 1 hour on a sunny summer day in Finland [6].

The UV range of the electromagnetic radiation spectrum extends from 10 nm to 400 and a major part of solar UV radiation which reaches the earth’s surface consists primarily of UVA radiation (90–99%) with the minor component of UVB radiation (1– 10%) nm.

Thereby, UV spectrum as shown is separated into four parts: UVA (315 nm to 400 nm), UVB (280 nm to 315 nm), UVC (200 nm to 280 nm) and UV Vacuum (100 nm to 200 nm) [5]. Decreasing wavelengths correspond with higher frequency radiation and a higher amount of energy per photon.

Recently published studies have demonstrated that UVA radiation can modulate a variety of biochemical processes, some of which are involved in the malignant transformation of skin [7,8] and mutagenesis [9,10].

UVA is known to cause severe oxidative damage via reactive oxygen species (ROS) [11], which can damage lipids [12], DNA [13] and induce apoptosis [14,15]. UVA may also play a significant role in the induction and development non-melanoma and melanoma skin cancers [10, 16,,17, 18,19].

UV light is generally used to disinfect environments [20] and in some cases for research purposes, such as in our laboratory. So, it is important to establish safety norms and standards in environments where there is ultraviolet radiation.

Therefore, to find out if chronic exposure to laboratory UV can cause health risks, we measured the irradiance inside and outside two types of laminar flows; with UV filter (Figure 1A) and other models which have a UV filter, but part of the glass is permanently open, which allows UV radiation to escape (Figure 1B) (Table 1).

**Table 1.** Laminar flow with radiation filter

| **A** | **Wavelengths** | **Inside (μW)** | **outside the glass(μW)** | **Outside 1m distance (μW)** |
| --- | --- | --- | --- | --- |
|  | **UVC 250nm** | 21.3/26.3/26.2/52.8 | 0.0/0.0/0.0/0.0 | 0.0/0.0/0.0/0.0 |
|  | **UVB 280nm** | 64.1/36.5/27.5/17.3 | 0.0/0.0/0.0/0.0 | 0/0/0.0/0.0/0.0 |
|  | **UVA 365nm** | 14.1/23.0/36.1/42.8 | 3.8/2.9/3.3/3.4 | 0.72/0.75/0.73/0.73 |

**Laminar flow without radiation filter**
| **B** | **Wavelengths** | **Inside (μW)** | **outside the glass(μW)** | **Outside 1m distance (μW)** |
| --- | --- | --- | --- | --- |
|  | **UVC 250nm** | 51.1/ 60.1/65.0/58.0 | 4.01/9.28/6.06/8.2 | 7.6/3.5/3.6/3.7 |
|  | **UVB 280nm** | 34.2/73.3/ 72.7/ 58.8 | 9.8/7.8/9.2/4.8 | 3.7/3.6/4.1/3.8 |
|  | **UVA 365nm** | 20.7/40.4/54.2/61.5 | 5.6/3.92/4.1/4.62 | 2.1/2.8/2.3/2.4 |

**Figure 1.**
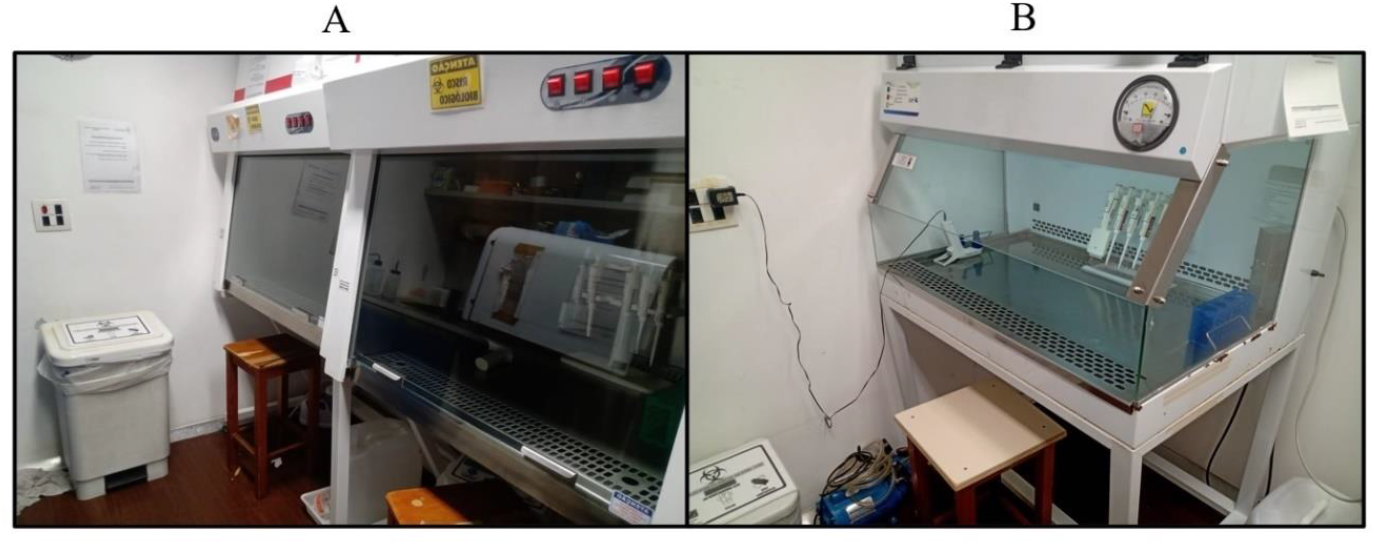

## Material and method

### Germicidal lamp

UV-C 30W tubular ultraviolet germicidal lamp for disinfection and sterilization.30W GERMICIDE lamp. Power: 30W. Model: TUV-30W PHILIPS. Base: G13. Bulb: T8. Total Length: 908.8 (max) mm. Diameter: 28 (max) mm. Main application: disinfection. Useful Life: 9000 h. Brand: Philips. Emits wave UV radiation with a peak of 253.7 nm (UV-C) with germicidal action.

### Laminar flows

The radiation dosage measurements were made in two types of laminar flow, equipped with the same type of UV lamp. A model with a UV filter on the glass, from the brand Pachane^®^ Pa 610, lamp height 52 centimeters (Figure 1 A). And a model without UV filter from the SOLUFIL^®^, model 288, lamp height 46 centimeters (Figure 1B).

### Mathematical formula to calculate the radiation dosage

To calculate the UV radiation dosage as a function of time, the following mathematical formula was used:

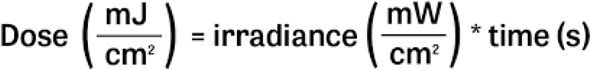

### Spectrum detector

Firstly, all environmental lights were turned off, only the internal UV of the flow remained on. To measure the wavelengths UVC 250nm, UVB 280nm and UVA 365nm was used a spectrum detector FieldMate Laser Power Meter. COHERENT (Figure 2) below.

**Figure 2.**
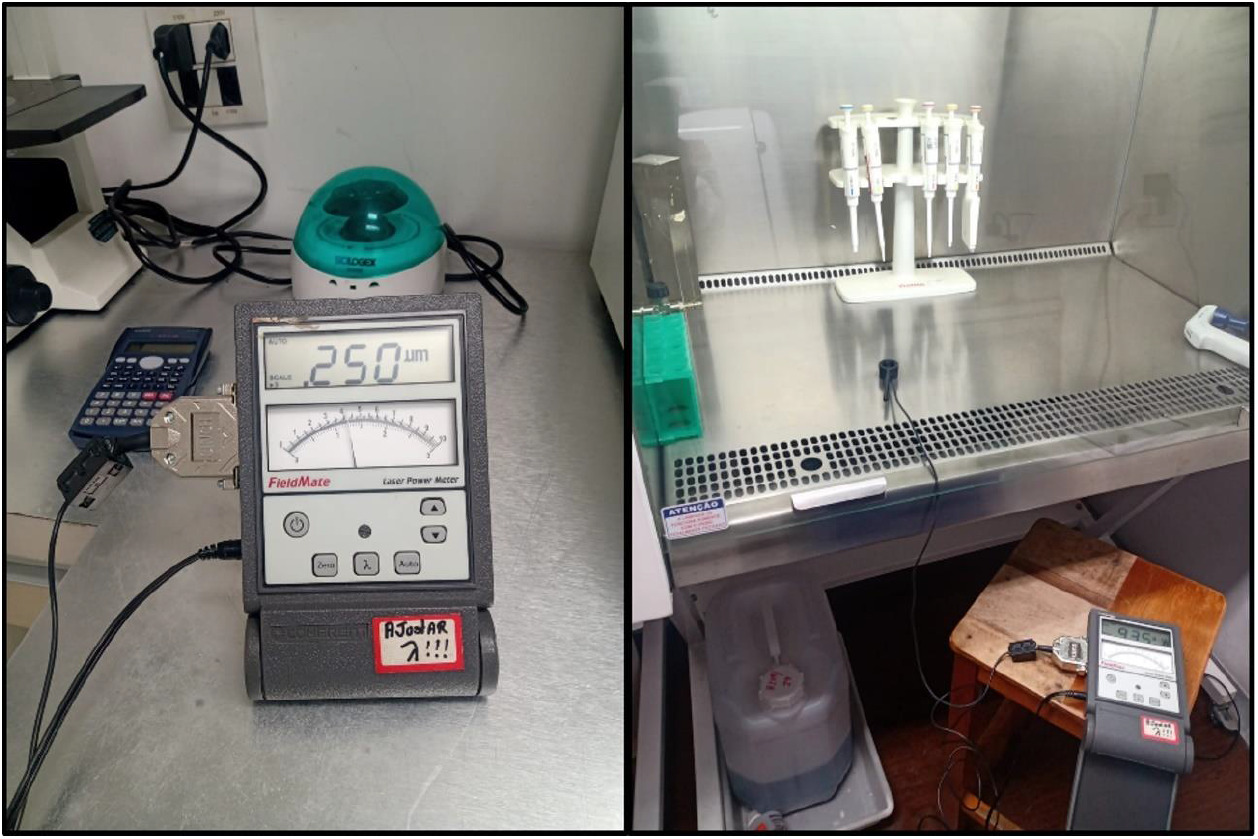

## Result and Discussion

The laminar flow, which has glass with UV filter, is efficient in reducing the UVC and UVB spectrum radiation that escapes into the laboratory environment (Figure 1 A) and (Table 1 A). For example, irradiance at a distance of one meter of UVC had an average of 0.73µW in UVA (Table 1 A), while at the same distance from the flow without UV filter (Figure 1 B), average measurements were UVC 4.6 µW, UVB 3.8 µW and UVA 2.4 µW (Table 1B). However, differences in irradiance between the lamps of the two laminar flows can be attributed to the time of use of the lamp and the height of lamp on each device, contributes to the scattering of radiation (Table 1).

For example, exposure without security protection for one hour a day and for 150 days a year in an environment with UV laboratory light, can lead to a dose of 27 J/cm^2^ of UVC, 30 J/cm^2^ of UVB and 15 J/cm^2^ of UVA (Table 1B).

These doses in specific wavelengths do not appear to high however the UV range of the electromagnetic radiation spectrum extends from 10 nm to 400 nm, so UV lights has health risk (table 2) [7].

**Table 2.**

| Band | Wavelength | Primary visual hazard | Other visual hazard | Other hazards |
| --- | --- | --- | --- | --- |
| UVA | 315 - 400 nm | Cataracts of Lens |  | Skin Cancer; Retinal Burns |
| UVB | 280 - 315 nm | Corneal Injuries | Cataracts of Lens;<br>Photokeratitis | Erythema; Skin Cancer |
| UVC | 100 - 280 nm | Corneal Injuries | Photokeratitis | Erythema; Skin Cancer |

In older laminar flows, see (Figure 1B), which do not have a safety system, it is possible the UV light to remain on, without the researcher noticing, so the doses received would be quite high, see measurements outside the glass in (Table 1B).

Depending on the wavelength and time of exposure, UV radiation may cause harm to the eyes and skin (table 2) [5,7,8,21].

In (figure 3) we see that UV lights occupy a band of the invisible electromagnetic field, with wavelength characteristics capable of interacting even with cellular genetic material.

**Figure 3.**
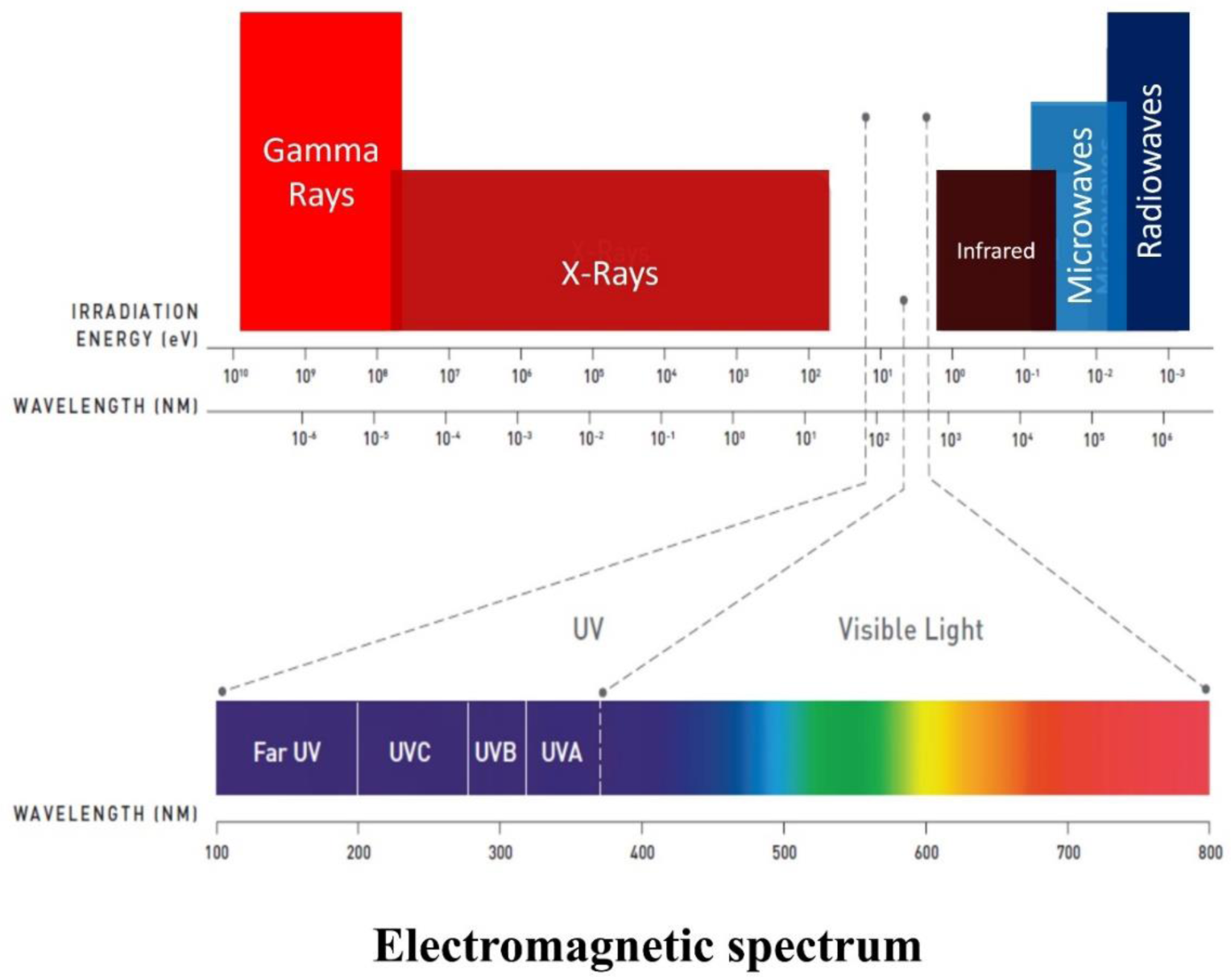

Regarding the biological effect of UV light, for example UVB has often been noted for its harmful effects on human skin, however each of the UV bands: UVA, UVB and UVC have a potential for damage (Table 2). Adverse health effects that may occur include erythema, photokeratitis, retinal burn, cataracts and others [21]. The (Table 2) summarizes these effects.

The shorter UVC wavelengths are typically absorbed in atmosphere, and thus are thought to have less long-term damaging effects on human tissue, as shown in (Figure 3)[22, 23 ]. However, prolonged direct exposure to chronic doses (table 1) of UVC light has caused eye and skin damage [24 ]. Because of this, exposure precautions are recommended when using UVC light.

Thus, UV light has a known impact on human tissue for example the scattering increases with decreasing wavelength (Figure 4) [25, 26, 27].

**Figure 4.**
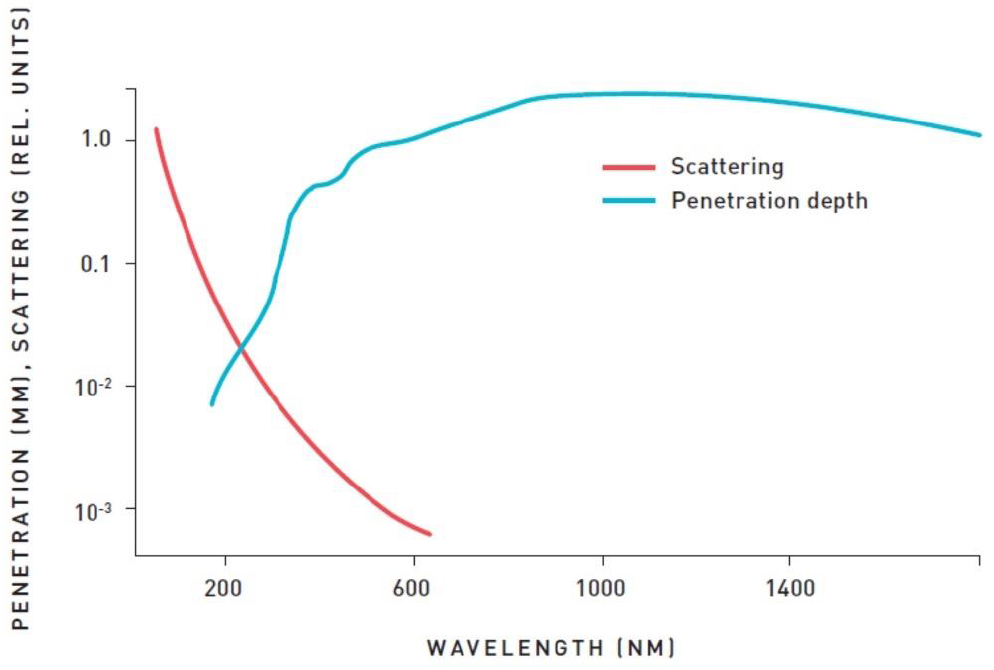
Impact of UV light on human tissue. In blue it is represented the penetration spectrum of light and UV radiation into human tissue. In red is represented the scattering increases with decreasing wavelength. Relative units Y axis and wavelength X axis. Nanometers (NM).

For example, acute exposure at high levels of UVC include redness or ulceration of the skin. For chronic exposures, there is also a cumulative risk, which depends on the amount of exposure at doses measured in (table 2). Thus, the long-term risk includes premature aging of the skin and skin cancer [28].

Therefore, even a few minutes’ exposure to the UVC radiation, at the doses found in (table 1), can result in photokeratisis and conjunctivitis. Both conditions through repetitive damage can cause premature aging of ocular structures, cataracts and blindness (table 2). [26, 29]. For example, Zuclich (1989) reported acute cataract induction by exposure to 337nm laser, in small dose of 1 J cm^2^ [30]. Like this, ultraviolet light can accelerate diseases linked to the aging process [31] .

Thus, it is important to use Personal Protective Equipment (PPE). UV radiation is easily absorbed by clothing, plastic or glass. Once absorbed, UV radiation is no longer active. When working with UV radiation during maintenance, service or other situations, personal protective equipment covering all exposed areas is recommended [32].

## Conclusions

The difficulty of organizing the arrangement of laminar flows in small spaces is common. Furthermore, laboratories have the challenge of maintaining overcrowded environments with researchers and employees.

Thus, personal safety training is important so personnel working with UVC fixtures or near UVC installations should be provided with training on health and safety topics, handling and maintenance of UVC sources, and first aid response after exposure to UVC light.

It is highly recommended that warning signs be placed where the UVC LEDs are electrically connected to warn incidental observers of the potential UV light exposure. These labels should be localized in all relevant languages and indicate that eye and skin hazard is probable and only authorized operators are permitted in the area.

Even in small spaces, intelligent organization of UV sources can be carried out, like this we can see in the (figure 5) three ways of organizing laminar flows: contraposed, juxtaposed or tandem respectively. It is important that areas in front of UV light have walls or panels that can absorb the radiation, and that there is no movement of people during the time that UV is on.

**Figure 5.**
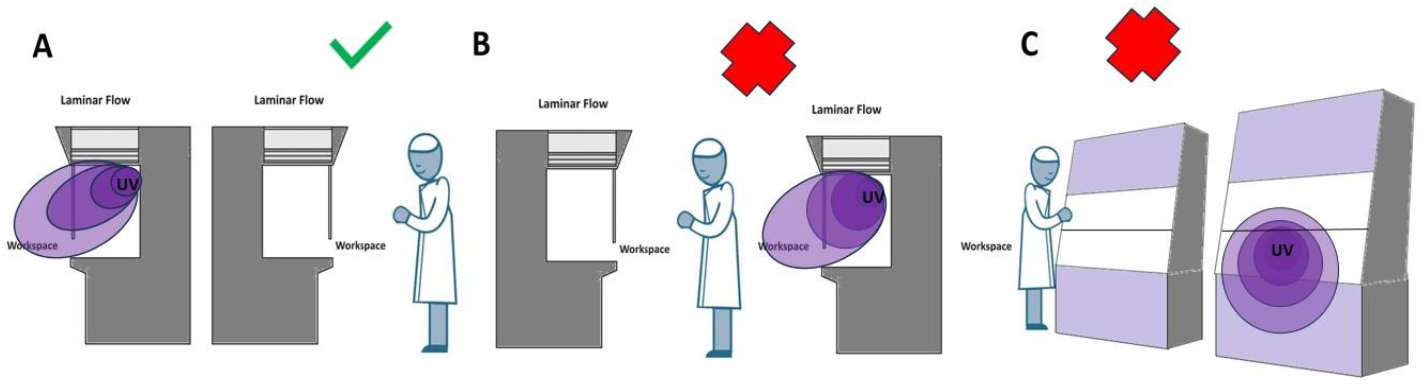
Ways of organizing laminar flows: (A) contraposed, (B) juxtaposed or (C) tandem. (A) is the correct way to organize UV sources. (B-C) are very common but incorrect ways as they expose other people to UV sources.

In adjacent areas warning labels should be placed outside access panels and doors to the UVC source as well as panels or doors where UVC radiation may penetrate or be reflected.

This way, UVC exposure can be reduced adopting the contraposed arrangement and through product safety controls (figure 5). For example, safety switches wired in series allow UVC sources to be turned off without exposing workers to UV light. Or placing ON/OFF switches for UVC light sources separate from general room lighting in locations only accessible by authorized persons. Switch locations should be locked or password protected to ensure that the UVC source is not accidentally turned on. Each UVC system should have the option of a viewport so workers can view the lamp assembly without the possibility of over-exposure to UVC.

Proper installation, monitoring, education of maintenance personnel, signage and use of safety switches can help to avoid overexposure. The operating instructions and recommendations for proper use of any UV system should be kept for reference to reduce hazardous exposure. These should be clearly visible for the operators or maintenance personnel specified by the system design to ensure safe operation.

Although, the effects of acute exposure to UV radiation are delayed. In the event of UV exposure, the following actions are recommended: See an ophthalmologist if eye damage is suspected; Treat skin lesions immediately; Follow Environment, Health and Safety (EHS) in your organization’s, to incident reporting procedure. These often require documentation of the date and time of incident, persons involved, equipment involved and type of injury.

Finally, when working around UVC devices, employees should use UV goggles and/or full face shields. Prescription glasses and normal safety glasses do not protect eyes from UV exposure, so equipment such as ANSI Z87 rated eyeglasses with wrap around lens to protect the side exposure is recommended. Cover any exposed skin using lab coats, nitrile gloves or other lab attire.

## Compliance with ethical standards Acknowledgments

I would like to thank my Professor Maurício da Silva Baptista for his confidence in my work. I would also like to thank the São Paulo Government Research Support Foundation (FAPESP) for funding.

